# Modular evolution and strain-specific partnerships: how *Hamiltonella defensa* shapes defense and symbiont communities in aphids

**DOI:** 10.1101/2025.10.31.685823

**Authors:** Nicole Lynn-Bell, Vilas Patel, Stephanie R. Weldon, Linyao Peng, Melissa Carpenter, Kyungsun Kim, Jacob A. Russell, Kerry M. Oliver

## Abstract

Insects frequently carry maternally-transmitted endosymbionts that mediate ecological interactions, including resource acquisition and host defense. All pea aphids (*Acyrthosiphon pisum*), for example, carry the obligate nutritional symbiont *Buchnera*, while most have one or more of the seven heritable facultative symbionts, which play diverse roles. A common symbiont, *Hamiltonella defensa*, confers protection against the parasitic wasp *Aphidius ervi* via a toxin-bearing bacteriophage called APSE, with strain-level variation in protection best predicted by APSE variant. Yet, little is known about *Hamiltonella*/APSE strain variation in the field, resulting in an incomplete understanding of the full arsenal of symbiont defenses and how these change over space and time. Here, we characterized *Hamiltonella*/APSE diversity in over 3,000 field-collected aphids from two North American populations across multiple years. We identified bacterial strains representing five *Hamiltonella* clades, seven APSE variants, and numerous toxin alleles, resulting in at least 38 distinct combinations. We found that APSEs moved laterally among *Hamiltonella* strains more readily than toxins among phage backbones, together generating modular defensive diversity. *Hamiltonella* strains exhibited strain-specific coinfection preferences with other symbionts, particularly *Fukatsuia*, *Rickettsia*, and *Rickettsiella*, indicating strain-level structuring of heritable bacterial communities. Geographic and temporal analyses revealed dramatic regional differences and rapid population turnover, with combinations conferring intermediate laboratory protection dominating despite the decline of highly protective strains, suggesting ecological success goes beyond parasitoid resistance alone. This extensive cryptic diversity provides standing genetic variation enabling rapid evolutionary responses to biological control efforts and anthropogenic stressors, including climate change, with implications for pest management and host adaptation.

## Introduction

Microbial symbionts represent one of the most pervasive yet underappreciated forces shaping terrestrial ecosystems, with bacterial partners estimated to infect most arthropod species (Douglas 2015, Weinert et al. 2015). While typically inconspicuous, microbial symbionts profoundly influence their hosts’ biology, from nutrient acquisition to defense against natural enemies (Douglas 2009, Hansen and Moran 2014, Oliver and Martinez 2014, Flórez et al. 2015).

Most insects carry environmentally acquired microbes that colonize exposed surfaces like the gut, as well as maternally transmitted endosymbionts that live within the body cavity and pass from mother to offspring across generations (Duron et al. 2008, Engel and Moran 2013, Kucuk 2020). Among these heritable endosymbionts, obligate nutritional partners have coevolved with hosts specialized on nutrient-poor diets such as plant sap, becoming essential for survival and reproduction (Baumann 2005). More widespread are heritable facultative symbionts, which, while not always essential, provide conditional benefits or manipulate host reproduction in ways that facilitate their persistence in host populations (O’Neill et al. 1997, Oliver et al. 2010).

The ecological success of facultative symbionts stems partly from their genetic flexibility. Unlike obligate symbionts, which show limited genetic variation due to ancient vertical transmission and gene loss (Chong et al. 2019), facultative symbionts typically maintain relatively large, dynamic genomes with extensive mobile genetic elements that enable adaptation to diverse hosts and tissues (Klasson et al. 2008, Moran et al. 2008, Penz et al. 2012). This genomic plasticity underlies functionally important strain-level variation within symbiont species. For instance, the globally distributed *Wolbachia* encompasses 17 putative supergroups, with some strains restricted to particular host taxa while others infect diverse arthropod species (Augustinos et al. 2011, Kaur et al. 2021). Critically, strains from the same supergroup can confer different phenotypes, while strains from different groups may provide similar benefits, highlighting the importance of understanding variation at the strain level rather than simply species presence or absence.

Aphids provide an exceptional system for investigating strain-level variation in heritable symbionts and its ecological consequences (Kucuk et al. 2025). While most aphid species universally harbor the obligate symbiont *Buchnera*, they also commonly carry one or more of nine documented symbiont species, though infection patterns vary significantly across taxa and populations (Oliver et al. 2010, Zytynska and Weisser 2016, Guo et al. 2017). Aphid facultative symbionts have become leading models of defensive symbiosis, demonstrating protection against viral and fungal pathogens, parasitoids, and thermal stress (Oliver et al. 2005, Scarborough et al. 2005, Schmid et al. 2012, Łukasik et al. 2013, Parker et al. 2013, Asplen et al. 2014, McLean and Godfray 2015, Hopper et al. 2018, Leybourne et al. 2020, McLean et al. 2020, Higashi et al. 2023, Higashi et al. 2024). Importantly, symbiont communities in pea aphids are not random assemblages but exhibit structured coinfection patterns, with certain symbiont combinations significantly enriched or depleted relative to chance expectations, suggesting complex interactions among community members (Rock et al. 2018, Peng et al. 2023).

The pea aphid, *Acyrthosiphon pisum*, a species complex of at least ten biotypes associated with different leguminous hosts (Peccoud et al. 2009), offers particularly rich opportunities for symbiont research. In North American populations of the alfalfa (*Medicago*) biotype, seven symbiont species occur regularly, with *Hamiltonella defensa* (Yersiniaceae: Enterobacterales) typically being the most prevalent (Russell et al. 2013, Smith et al. 2015, Carpenter et al. 2021). This symbiont provides crucial protection against *Aphidius ervi* (Hymenoptera: Braconidae), the primary parasitoid of alfalfa-feeding pea aphids (Angalet and Fuester 1977, Oliver et al. 2003). Importantly, *Hamiltonella’s* protective efficacy depends critically on strain identity and associated bacteriophages called APSEs. Previous studies show that *Hamiltonella* isolates vary in bacterial and APSE genotypes which correspond to differing defensive capabilities (Degnan and Moran 2008, Oliver et al. 2009, Martinez et al. 2018b, Oliver and Higashi 2019, Patel et al. 2023). However, important gaps remain in our knowledge of the genomic and functional diversity of *Hamiltonella* present in natural populations, including whether the most protective strains are the most common in field populations.

The modular organization of defensive elements within the *Hamiltonella*-APSE system creates opportunities for generating novel protective phenotypes through recombination and horizontal transfer. Phylogenetic and laboratory studies demonstrate that APSE can move between bacterial lineages, while comparative genomics reveals evidence for toxin shuffling among phage backbones (Degnan and Moran 2008, Brandt et al. 2017, Lynn-Bell et al. 2019, Boyd et al. 2021, Patel et al. 2023), suggesting that natural populations may harbor extensive cryptic diversity created through recombination of defensive modules. However, the evolutionary constraints governing these processes remain poorly understood.

Field studies reveal that *Hamiltonella* infections fluctuate rapidly in response to parasitism pressure and environmental conditions (Ives et al. 2020, Carpenter et al. 2021, Gimmi and Vorburger 2021, Smith et al. 2021), yet our understanding of the strain-level diversity underlying these dynamics remains incomplete. Given the strong correlation between strain variation and host protection demonstrated in genomic studies, comprehensive characterization of *Hamiltonella* and APSE diversity in natural populations is essential for understanding how heritable microbiomes are maintained, and for appreciating the full defensive arsenal available to aphids in nature. Studies of this diversity have further, practical implications for biological control, as aphid populations may possess substantial standing genetic variation enabling rapid evolutionary responses to parasitoid introductions (Desneux et al. 2018, Vorburger 2018).

Here, we examined the heritable facultative symbiont community of over 3,000 pea aphids from two North American populations of the *Medicago sativa* biotype across multiple years. Using established multilocus sequence typing combined with novel diagnostic approaches, we provide the most comprehensive analysis of *Hamiltonella* strain diversity conducted to date. We address three key questions: (1) How extensive is cryptic diversity within *Hamiltonella* and APSE? (2) Do specific *Hamiltonella* strains show preferential associations with other symbionts, indicating a role in structuring microbial communities (Doremus and Oliver 2017, Peng et al. 2023)? (3)

How does strain-level diversity vary across space and time? This work reveals the hidden complexity driving symbiont-host dynamics in nature and demonstrates how strain-level variation shapes both defensive function and community structure.

## Methods

### Aphid Sample Collection

Pea aphids were collected from alfalfa (*Medicago sativa*) fields across two geographic regions over multiple years: 13 fields in Arlington, Wisconsin (2011-2019) and 7 fields in Fargo, North Dakota (2015-2018). Wisconsin field sites included locations 218, 219, 246, 301, 310, 349, 412, 506SE, 591, 593, 576N, 744, and N1902, while North Dakota sites were designated B, R, Y, 19, 25, 26, and 28. To minimize pseudoreplication from clonal reproduction, individual aphids were collected at least 10 meters apart within each field. Collections occurred early in the growing season (late May to early June) when pea aphid populations first became abundant.

## Symbiont Community Characterization

### General Screening

Field-collected aphids were transported to the laboratory where parthenogenetic adults were isolated individually and allowed to reproduce to establish clonal lines. DNA was extracted from the original field-collected mothers using the E.Z.N.A Tissue DNA Kit (Omega Biotek) following the whole-aphid protocol and stored at -80°C until analysis.

For samples collected from 2014-2019 (n = 2,232), we screened individual aphids for seven facultative symbionts known to occur in pea aphids: *Hamiltonella*, *Rickettsia*, *Fukatsuia*, *Rickettsiella*, *Regiella*, *Serratia*, and *Spiroplasma*. We used established PCR primers and protocols described in Russell et al. (2013) (Table S1). To detect unexpected symbionts that might have colonized the study populations, we performed Illumina MiSeq amplicon sequencing targeting the hypervariable V3-V4 regions of bacterial 16S rRNA using primers V3f (5’- CCTACGGGAGGCAGCAG-3’) and V4r (5’-GGACTACHVGGGTWTCTAAT-3’) following protocols described by (Kozich et al. 2013). Pooled samples (∼100 aphids per geographic location) were sequenced across multiple years with appropriate negative controls. This analysis confirmed only the seven expected symbiont species were present, so no additional symbionts were included in subsequent analyses.

### *Hamiltonella* Strain and APSE Characterization

For all *Hamiltonella*-positive aphids, we characterized bacterial strain identity, associated APSE (*<u>A</u>. <u>p</u>isum* <u>s</u>econdary <u>e</u>ndosymbiont) bacteriophage variants, and toxin composition. Traditional Sanger sequencing of specific housekeeping genes (*ptsI*, *recJ*, *accD*, *murE*, and *hrpA*) distinguished *Hamiltonella* strains using a multilocus sequence typing (MLST)-like approach (Degnan and Moran 2008). However, since typing loci among strains vary only by a few SNPs, they cannot be easily adapted for rapid strain diagnostics using PCR. To enable rapid strain identification, we developed insertion sequence (IS)-based diagnostic PCR assays using five primer sets (Table S1) that specifically amplify different *Hamiltonella* clades (A-E) based on unique IS element distributions along the main chromosome.

To assess accuracy of our IS-based approach, we Sanger sequenced two MLST loci (*recJ* and *ptsI*) from 95 samples and compared results with IS-based clade assignments. For samples that failed to amplify with IS-based primers, we reverted to traditional MLST approaches to capture potentially novel bacterial genotypes.

APSE variants were characterized by Sanger sequencing six gene fragments (P3, P35, P41, P38, P45 and P51) spanning the three APSE modules that encode structural and regulatory functions (Degnan and Moran 2008, Boyd et al. 2021). Toxin identity was determined using diagnostic PCR with primers targeting known APSE-encoded virulence factors (Table S1). For samples potentially harboring novel toxin alleles, we performed Sanger sequencing of the relevant gene fragments. The two most common APSE variants (APSE2 and APSE8) encode highly similar *cdtB* toxin alleles that we distinguished using either: (1) Sanger sequencing of *cdtB* gene fragments, or (2) restriction digest analysis with TscAI endonuclease following FastDigest protocols (Thermo Scientific, Marietta, OH) to differentiate cdtB1 (APSE2- associated) from cdtB2 (APSE8-associated).

### Phylogenetic and Diversity Analyses

*Hamiltonella* and APSE genotypes were considered distinct when they exhibited sequence variation at any amplified typing locus. *Hamiltonella*-APSE combinations were treated as distinct genotypes when: (1) identical *Hamiltonella* strains carried different APSE variants, or (2) identical APSE backbones encoded different toxin variants.

All sequences were aligned using MUSCLE (Edgar 2004) in Geneious Prime https://www.geneious.com). Recombination analysis using RDP4 (Martin et al. 2015) identified recombination signals in two APSE loci (P41 and P51). Consequently, we limited our analysis to four loci (P3, P35, P38, P45) spanning the three non-virulence modules for phylogenetic analyses.

Phylogenetic trees were constructed for both individual MLST genes and concatenated sequences. We used ModelFinder (Kalyaanamoorthy et al. 2017) to identify the best substitution models for individual loci with partitioned models for concatenated sequences. We estimated maximum likelihood phylogenies using IQ-TREE webserver with 10,000 ultrafast bootstrap replicates to assess branch support (Trifinopoulos et al. 2016) and conducted network analyses using SplitsTrees version 6 (Huson and Bryant 2024). Individual locus alignments were analyzed separately to identify conflicting phylogenetic signals, and a concatenated alignment of all five loci was subjected to network analysis using the Neighbor-Net algorithm with default parameters.

### Statistical analyses

All analyses were conducted using R Statistical Software (v4.5.1; R Core Team, 2025, Vienna, https://www.R-project.org/) with the RStudio interface (Posit team, 2025, Boston [http://www.posit.co/]).

### Symbiont Occurrence Patterns

To test whether particular symbiont infections occurred more or less frequently than expected by chance, we performed pairwise Fisher’s exact tests and binomial tests (R packages fisher.test and bionm.test). Multi-way infections were collapsed into pairwise categories to focus on biologically meaningful interactions while reducing multiple comparison issues. For each symbiont pair, we constructed 2×2 contingency tables comparing observed versus expected coinfection frequencies under statistical independence. Expected coinfection frequencies were calculated as the product of individual symbiont prevalences multiplied by total sample size. We applied Benjamini-Hochberg false discovery rate (FDR) correction (p.adjust package with method = “fdr”) across the 15 pairwise tests, with associations considered significant at FDR-adjusted p < 0.05.

Associations between 1) *Hamiltonella* clades and APSE variants, and 2) toxin types and APSE variants, were analyzed using chi-square tests of independence. Standardized residuals identified specific significant associations, and FDR corrections controlled for multiple testing. Shannon diversity indices (vegan package) were calculated for each clade to quantify APSE type richness and evenness.

Predictive relationships were assessed using multinomial logistic regression models using the nnet package. Model 1 predicted APSE variant from *Hamiltonella* clade, while Model 2 predicted toxin type from APSE variant. Model performance was evaluated using pseudo R² (McFadden’s), overall accuracy, confusion matrices (caret package), and odds ratios with 95% confidence intervals. ANOVA tests (car package) assessed overall model significance. Model outputs were processed using the broom package for tidy formatting.

We analyzed associations between *Hamiltonella* clades and coinfecting bacterial symbionts using Fisher’s exact tests with Monte Carlo simulation (10,000 replicates) due to computational limitations with large contingency tables.

First, we assessed associations between *Hamiltonella* clades (A-E) and the most common coinfecting symbionts *Fukatsuia*, *Rickettsiella*, and *Rickettsia* (N = 468, 131, 129, respectively) using 6×2 Fisher’s exact tests for each symbiont across all *Hamiltonella* clades plus single infections, with FDR correction applied across the three symbiont tests.

Second, we tested for non-random associations between APSE variants and coinfecting symbionts using each of four symbiont taxa (*Regiella*, *Fukatsuia*, *Rickettsiella*, and *Rickettsia*) across five APSE variants (APSE1, APSE2, APSE3, APSE8, and APSE11) found in coinfections. APSE9 was excluded due to absence of symbiont coinfections. For each symbiont, 2×5 contingency tables were constructed comparing focal symbiont counts versus all other symbionts across APSE variants.

Third, we conducted Fisher’s exact tests for three symbiont taxa (*Fukatsuia*, *Rickettsiella*, and *Rickettsia*) associated with B-clade *Hamiltonella* across its five phage variants (B1, B2, B3, B8, and B10). For both APSE and B-phage analyses, 2×5 contingency tables were constructed for each symbiont, and FDR correction was applied across tests within each analysis. Chi-square tests of independence assessed overall associations between phage variants and symbiont community composition for both APSE and B-phage datasets.

### Multiple *Hamiltonella* Coinfections

To test whether infections with multiple *Hamiltonella* strains occurred at frequencies expected from random pairing based on clade abundance, we calculated expected coinfection frequencies for all possible clade pairs using the product of their marginal frequencies. We performed chi-square goodness-of-fit tests comparing observed versus expected frequencies for each combination with sufficient expected counts (≥1). P-values were adjusted for multiple comparisons using FDR. An overall multinomial goodness-of-fit test assessed whether the entire coinfection pattern deviated significantly from random expectation.

### Spatial Diversity Analyses

Symbiont diversity at the species level as well as *Hamiltonella* clade (A - E) and APSE variant (1, 2, 3, 8, 9, 10, 11) diversity and combinations (A2, B1, etc.) were calculated using the Shannon diversity index (H’) to assess both species richness and evenness across collection sites. Shannon diversity was computed using the diversity function in the vegan package (v2.6-4).

Additional diversity metrics including Simpson’s diversity index and Pielou’s evenness (J’) were calculated for comparative analysis. Statistical differences in Shannon diversity between Wisconsin (WI) and North Dakota (ND) collection sites were tested using Hutcheson’s t-test, which accounts for unequal sample sizes and variance differences between sites. Compositional differences in combination distributions between Wisconsin and North Dakota were evaluated using chi-square tests of independence, with standardized residuals identifying specific combinations driving compositional differences. Bray-Curtis dissimilarity quantified overall community compositional differences between sites.

Total richness of *Hamiltonella* + APSE combinations was estimated using the Chao1 non-parametric estimator, which accounts for unseen species based on the frequency of rare combinations (singletons and doubletons). The Chao1 estimator was calculated using the estimateR function in the vegan (v2.6-4) package (https://cran.r-project.org/package=vegan). Sampling completeness was assessed as the ratio of observed to estimated richness, and 95% confidence intervals were calculated using standard errors. Site-specific Chao1 estimates were computed separately for North Dakota and Wisconsin to evaluate regional completeness. Theoretical maximum richness was calculated as the product of observed *Hamiltonella* clades (5) and APSE types (7), totaling 35 possible combinations.

### Temporal Changes in Symbiont Community Structure

Temporal changes in symbiont community structure were analyzed at two levels: symbiont community composition and *Hamiltonella*-phage combinations. GLMMs were fitted using the lme4 package (Bates et al. 2015) with Wald tests performed using the car package (https://www.john-fox.ca/Companion/.). Diversity metrics and multivariate analyses used the vegan package (v2.6-4). Compositional data analysis was performed using the compositions package, data manipulation used dplyr and tidyr, with visualizations created using ggplot2.

### *Hamiltonella* + phage combination analysis

Data consisted of counts of 17 different *Hamiltonella* clade + phage combinations sampled from Wisconsin populations across six years (2014-2019), with sample sizes ranging from 34 to 160 individuals per year (total n = 635). For individual combination analyses, we fitted binomial GLMMs with logit links, modeling the proportion of each combination relative to the total sample size per year. Year was included as a centered continuous fixed effect, and a random intercept for year was included to account for temporal autocorrelation. Models were fitted using maximum likelihood with the bobyqa optimizer. We focused analyses on four major combinations (A2, B1, B8, D3) that had sufficient sample sizes (≥5 total observations) for reliable estimation. Statistical significance of temporal trends was assessed using Type II Wald chi-square tests. Community-level changes were evaluated by calculating Shannon diversity, species richness, and total abundance for each year, with temporal trends assessed using linear regression.

## Symbiont community analysis

Data consisted of counts for seven symbiont categories (*Fukatsuia*, *Regiella*, *Rickettsiella*, *Serratia*, *Hamiltonella*, *Rickettsia*, and Uninfected) sampled from Wisconsin populations across the same six-year period, with sample sizes ranging from 115 to 986 individuals per year (total n = 2,595). Individual symbiont temporal trends were assessed using the same GLMM approach as described above.

Community-level diversity changes for both datasets were evaluated by calculating Shannon diversity, Simpson diversity, evenness, dominance, and species richness for each year, with temporal trends assessed using linear regression. Multivariate community analysis employed PERMANOVA (adonis2) to test for overall compositional changes over time using Bray-Curtis dissimilarity matrices, with year treated as a continuous predictor.

## Results

### Overview of symbiont infections

We screened 2232 pea aphids and recovered all seven previously reported facultative symbiont species (Figure 1). *Spiroplasma* was detected in fewer than 0.1% of aphids and was excluded from subsequent analyses due to its rarity.

**Figure 1:**
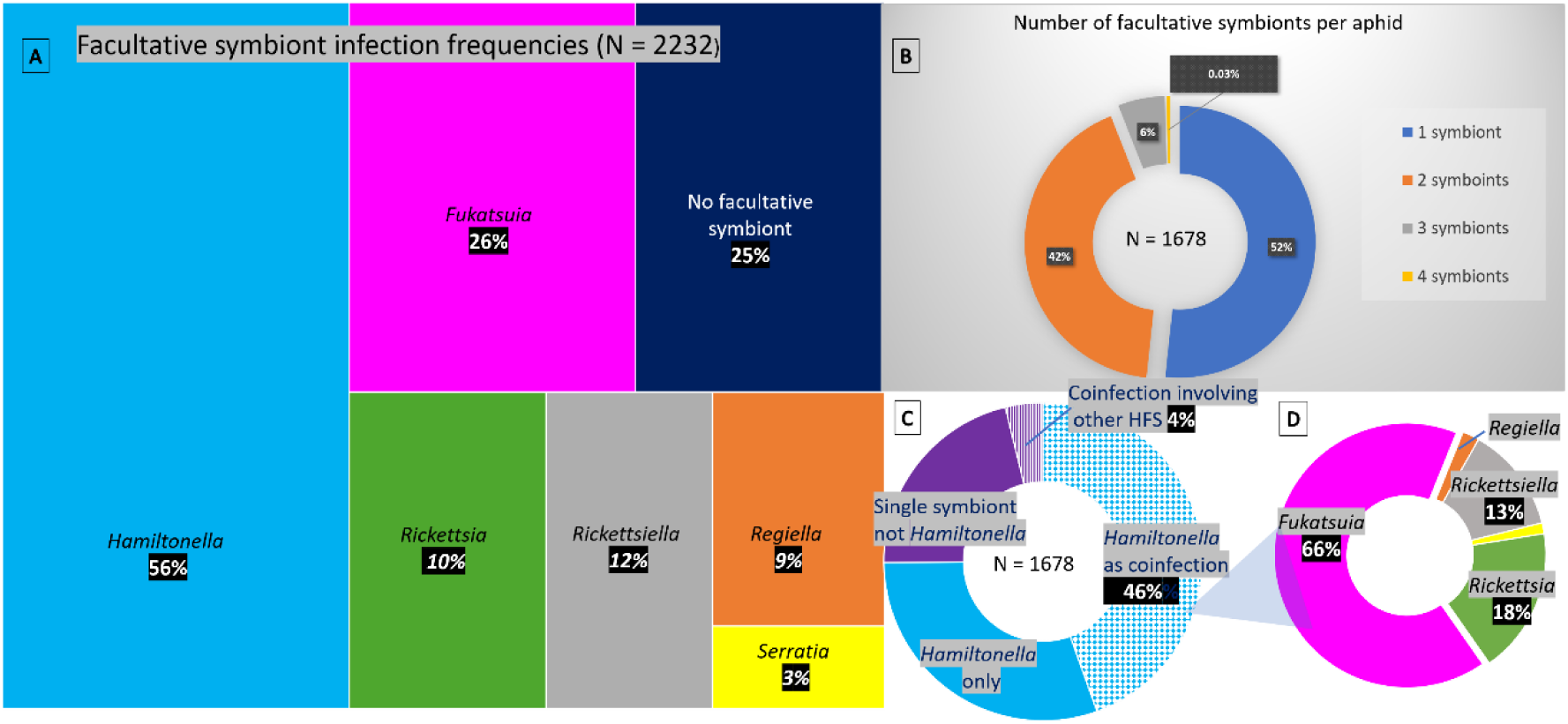
Facultative symbiont infection frequencies across sampled Wisconsin and North Dakota pea aphids feeding on alfalfa. A) Symbiont species infection frequencies, note that percentages total more than 100% due to multiple symbiont infections in individual aphids, B) number of facultative symbionts per aphid, C) indicates percentages of *Hamiltonella* single infections, total of other symbiont single infections, coinfections involving *Hamiltonella,* and coinfections involving other symbionts, d) other symbionts forming coinfections with *Hamiltonella*

### Infection Patterns and Prevalence

Nearly one-quarter of pea aphids contained only the obligate symbiont *Buchnera*, while the majority carried one or more facultative symbionts (mean = 1.16; Figure 1A). Among aphids harboring facultative symbionts, the average was 1.6 symbionts per individual aphid (Figure 1B). Specifically, 39% of aphids carried a single facultative symbiont, 31% carried two, and about 5% carried three or four symbionts (Table 1). *Hamiltonella* was the most common facultative symbiont, occurring in 56% of sampled aphids. The remaining five facultative symbionts showed variable prevalence, ranging from 3% (*Serratia)* to 26% (*Fukatsuia*) (Figure 1A).

**Table 1:**
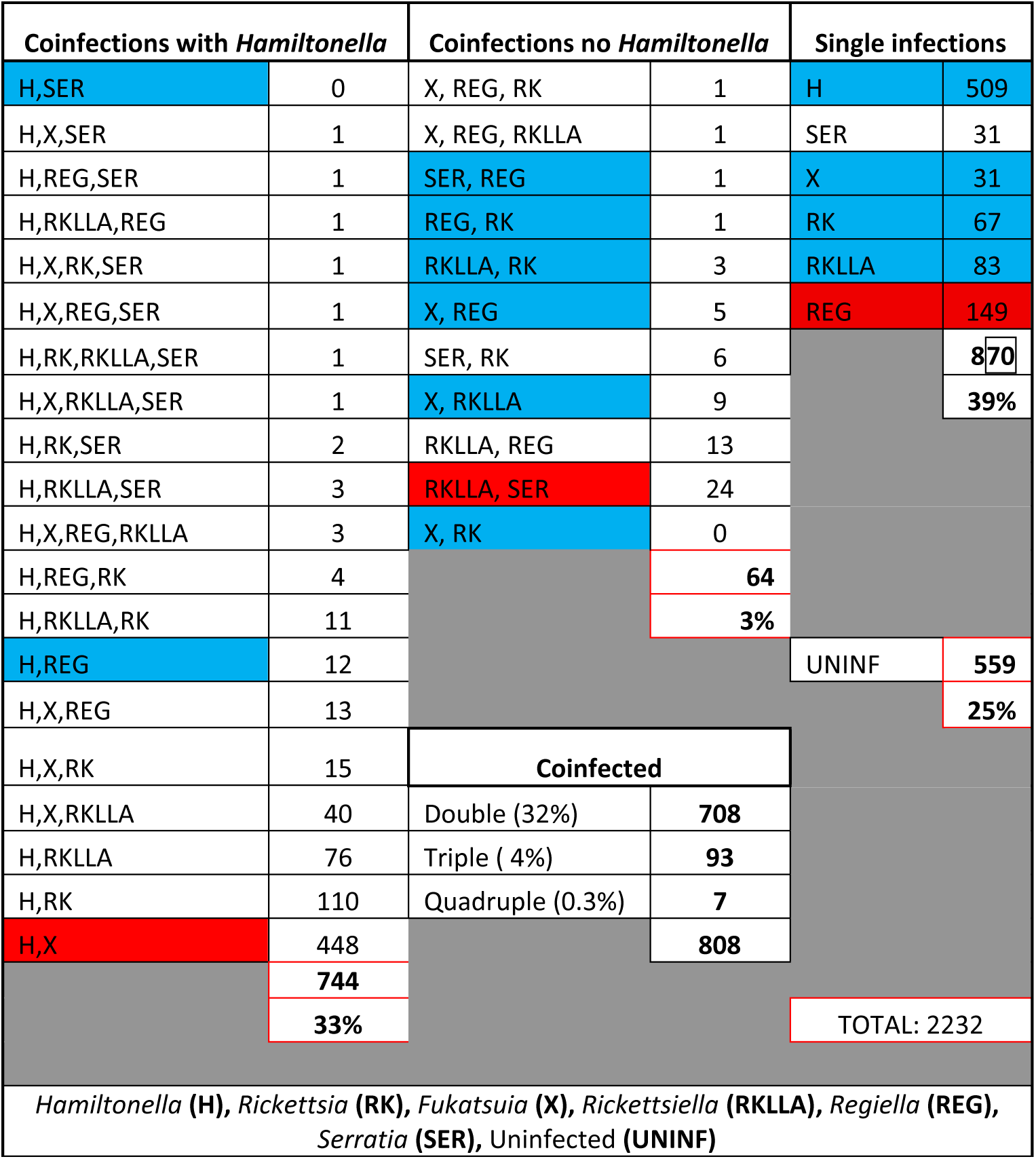
Summary of symbiont species types across pea aphid populations

| Coinfections with *Hamiltonella* |  | Coinfections no *Hamiltonella* |  | Single infections |  |
| --- | --- | --- | --- | --- | --- |
| H,SER | 0 | X, REG, RK | 1 | H | 509 |
| H,X,SER | 1 | X, REG, RKLLA | 1 | SER | 31 |
| H,REG,SER | 1 | SER, REG | 1 | X | 31 |
| H,RKLLA,REG | 1 | REG, RK | 1 | RK | 67 |
| H,X,RK,SER | 1 | RKLLA, RK | 3 | RKLLA | 83 |
| H,X,REG,SER | 1 | X, REG | 5 | REG | 149 |
| H,RK,RKLLA,SER | 1 | SER, RK | 6 |  | 870 |
| H,X,RKLLA,SER | 1 | X, RKLLA | 9 |  | 39% |
| H,RK,SER | 2 | RKLLA, REG | 13 |  |  |
| H,RKLLA,SER | 3 | RKLLA, SER | 24 |  |  |
| H,X,REG,RKLLA | 3 | X, RK | 0 |  |  |
| H,REG,RK | 4 |  | 64 |  |  |
| H,RKLLA,RK | 11 |  | 3% |  |  |
| H,REG | 12 |  |  | UNINF | 559 |
| H,X,REG | 13 |  |  |  | 25% |
| H,X,RK | 15 |  | Coinfected |  |  |
| H,X,RKLLA | 40 | Double (32%) | 708 |  |  |
| H,RKLLA | 76 | Triple ( 4%) | 93 |  |  |
| H,RK | 110 | Quadruple (0.3%) | 7 |  |  |
| H,X | 448 |  | 808 |  |  |
|  | 744 |  |  |  |  |
|  | 33% |  |  | TOTAL: 2232 |  |
| *Hamiltonella* (H), *Rickettsia* (RK), *Fukatsuia* (X), *Rickettsiella* (RKLLA), *Regiella* (REG), *Serratia* (SER), Uninfected (UNINF) |  |  |  |  |  |

### Coinfection Dynamics

While all symbiont species were observed in coinfections, several species displayed distinct association patterns. *Hamiltonella*, *Fukatsuia*, *Rickettsia* and *Rickettsiella* occurred in coinfections more often than expected by chance (Table 1). *Hamiltonella* showed a moderate tendency toward coinfection, occurring 1.46 times more frequently as a coinfection (59%) than as a single infection (41%) (one-tailed binomial test, n = 1,252; 95% CI: 0.571-1.000; FDR-adjusted *p* < 0.001) (Figure 1C; Table 1), while *Fukatsuia* was 17X more likely to occur as a coinfection than a single infection, doing so 95% of the time (binomial test, n = 570 95% CI 0.927, 1.000, FDR-adjusted p < 0.001). Specifically, *Fukatsuia* formed a coinfection with *Hamiltonella* 97% of the time, which was significantly more likely than with other symbionts (Table 1D; binomial test, n = 539 95% CI 0.955, 1.0000, p < 0.0001). In contrast, *Regiella* (72%; binomial test, n = 206 95% CI 0.667, 1.000, FDR-adjusted p < 0.001) was more likely than expected by chance to occur as single infections, while *Serratia* was not more likely to occur in single versus coinfections (binomial test, n = 73, 95% CI 0.575, 1.000, FDR-adjusted p =0.12)

In total, we found 35 total species-level symbiont combinations (Table 1). Eleven combinations were singletons. Some combinations, such as *Hamiltonella* and *Serratia* were never observed.

*Hamiltonella* participated in 19 different symbiont combinations representing 92% of observed coinfections (N = 808 aphids) (Table 1). Two symbiont combinations showed significant enrichment relative to chance expectations. *Hamiltonella* and *Fukatsuia* showed a very strong positive association, co-occurring 1.63 times more frequently than expected by chance (Table 1; OR = 14.19, FDR-adjusted p < 0.001), as did *Serratia* + *Rickettsiella* (OR 4.07; FDR-adjusted p < 0.001). Conversely, numerous combinations occurred significantly less frequently than expected, such as *Hamiltonella* + *Regiella* (OR = 0.13, FDR-adjusted p < 0.001) and *Hamiltonella* + *Serattia* (OR = 0.00, FDR-adjusted p < 0.001), the latter being a combination not observed. Significantly enriched (red) and depleted combinations (blue) are presented in Table 1.

### *Hamiltonella* and APSE diversity

As the most frequently sampled facultative symbiont in our study, and the species involved in the majority of coinfections, we examined *Hamiltonella* diversity in greater detail. Using sequence typing at five loci combined with network analyses and maximum likelihood phylogenetic reconstruction, we found that most isolates clustered within the previously established A-E clade classification scheme (Figure 2, Figure S1A) (Patel et al. 2023). Each isolate in Figure 2 represents a novel genotype and thus visually overrepresents diversity. Analysis of a representative subset of 109 *Hamiltonella* isolates across all five typing loci revealed that a large majority (85%) of isolates for a given clade exhibited identical alleles at each locus. For example, A-clade isolates typically displayed the AAAAA genotype (noting that the *recJ* locus overlaps with C-clade: uppermost A-clade isolate in Figure 2), while B-clade isolates showed BBBBB (Figure 2, noting that the *hrpA* locus overlaps with D clade). However, we also detected substantial within-clade variation, identifying at least 14 distinct bacterial genotypes. For instance, B-clade isolate 246-11 possessed a D-clade *ptsI* allele while maintaining typical B- clade alleles at the remaining four loci (Figure 2, Figure S1B).

**Figure 2:**
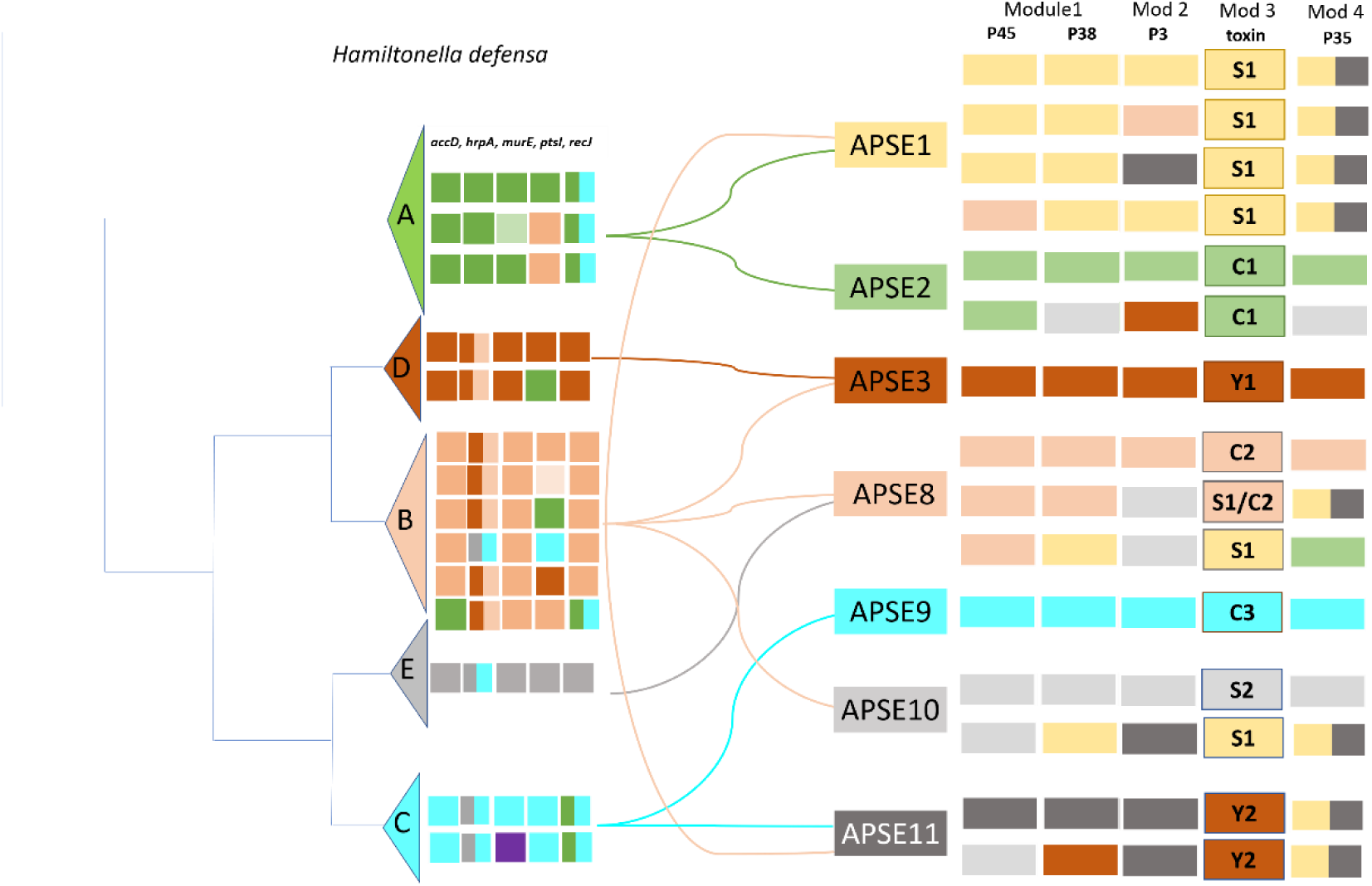
Genetic diversity and lineage associations among *Hamiltonella* and APSE isolates. A phylogenetic tree (left) illustrates the relationships among five bacterial clades (A–E) of *Hamiltonella*. Each row of colored boxes represents a single *Hamiltonella* isolate, with the color indicating the lineage of a specific typing locus (loci shown in the top inset). This analysis reveals distinct genetic patterns within the bacterial host. The seven *Hamiltonella* clades are associated with seven distinct APSE phage backbones (center, labeled APSE1-11). The colored boxes on the right represent the toxin genes found within each APSE backbone. Toxin alleles were categorized into three groups: Shiga-like toxin (S), cytolethal distending toxin (C), and YD-repeat toxin (Y), with different numbers indicating distinct alleles (e.g., S1, S2). This figure demonstrates the extensive co-evolutionary relationships and genetic variability observed between *Hamiltonella* clades and their associated APSE phages, highlighting the mosaic nature of the phage genomes and the diversity of their encoded toxins.

APSE variant analysis showed that most recovered sequences fell within previously characterized groups APSE1, 2, 3, 8, 9, 10 and 11 (Patel et al. 2023). However, we identified multiple novel haplotypes that require further characterization (Figure S1C). But as with *Hamiltonella*, among a subset of 134 APSE isolates examined across typing loci, most (72%) isolates shared a common set of co-inherited alleles for a given variant, while the remainder exhibited combinations associated with different APSE variants (compare the two APSE2s in Figure 2). Within the APSE virulence module, we identified seven distinct toxin alleles, each belonging to one of the three previously established toxin groups (Figure 2). Considering *Hamiltonella* and APSE phage backbone typing loci plus toxin alleles, we conservatively recovered at least 38 distinct combinations of *Hamiltonella*.

In addition to characterizing total diversity, we analyzed the relative frequencies of *Hamiltonella*, APSE, and primary toxins to assess their prevalence in our field collections. Among the 860 *Hamiltonella* isolates identified, B-clade strains were most abundant (59%), followed by A-clade (25%) and D-clade isolates (10%) (Figure 3A). C-clade *Hamiltonella* was very rare.

**Figure 3:**
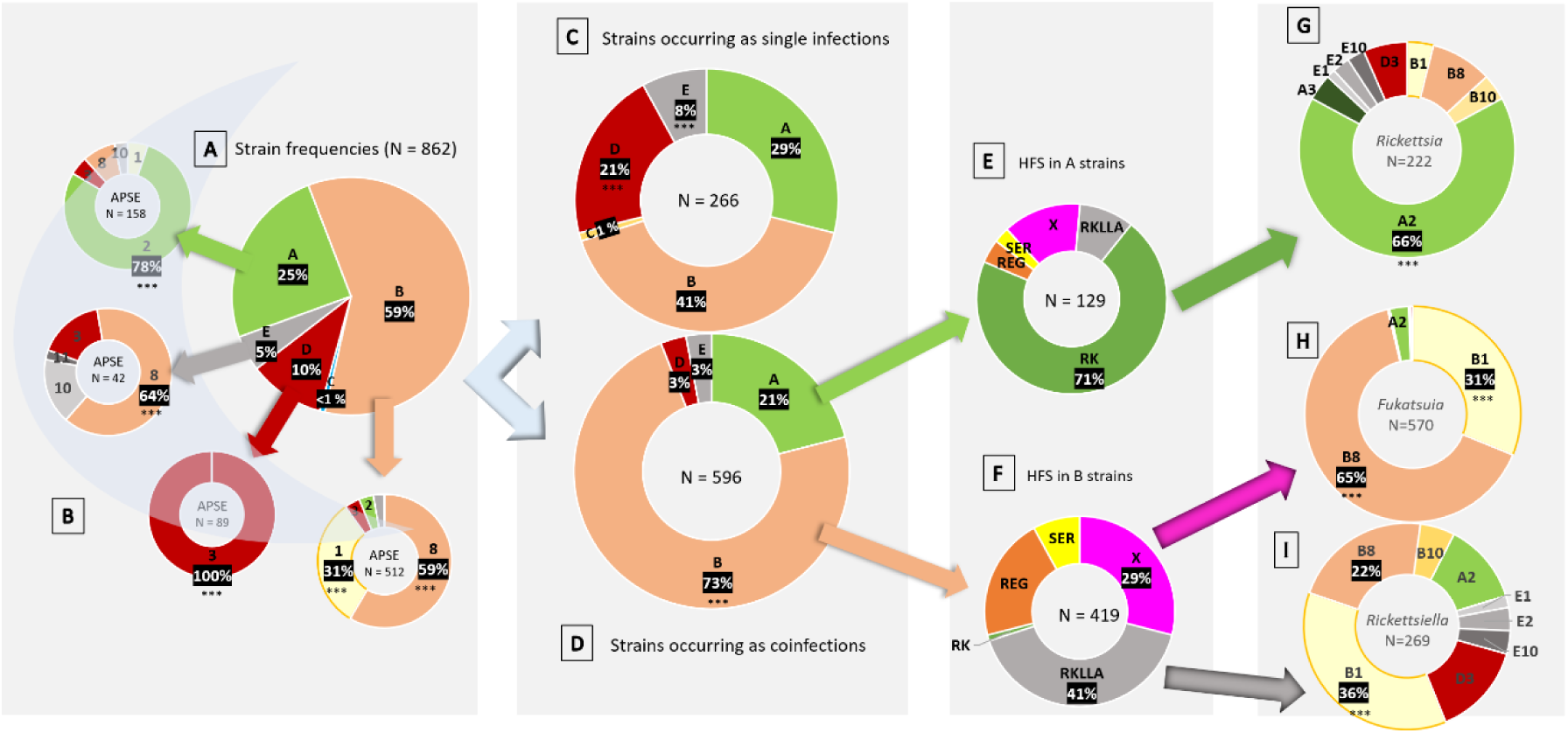
***Hamiltonella* and APSE infection patterns.** This figure details the prevalence and coinfection dynamics of *Hamiltonella* and its associated APSE phages across an aphid population. Panel A shows the overall frequency of the five major *Hamiltonella* clades (A–E) in the total aphid sample, while Panel B details the frequency of the most common APSE phage variants associated with each *Hamiltonella* clade. The figure then distinguishes between aphids with single-symbiont infections (Panel C) and those with coinfections (Panel D). Panels E and F break down the specific coinfecting symbionts associated with *Hamiltonella* clades A and B. These symbionts are abbreviated as follows: *Rickettsia* (RK), *Fukatsuia* (FUK), *Rickettsiella* (RKLLA), *Regiella* (REG), and *Serratia* (SER). Panels G, H, and I provide a more detailed look at the APSE-*Hamiltonella* combinations , namely those coinfected with *Rickettsia*, *Fukatsuia*, and *Rickettsiella*, respectively. Asterisks (***) denote a statistically significant difference in prevalence (p<0.001).

*Hamiltonella* clades exhibited highly significant non-random associations with APSE variants (Figure 3B χ² = 1878.14, df = 24, p < 0.001, Cramér’s V = 0.75), as indicated by multinomial logistic regression, which demonstrated moderate predictive power (pseudo R² = 0.40, accuracy = 66.9%). Despite this, clear evidence of repeated horizontal transfer emerged: isolates from four *Hamiltonella* clades harbored three or more distinct APSE variants, while only D-clade exhibited perfect specificity for APSE3 (100%, n = 89, Shannon H’ = 0)(Figure 3B). Notably, APSE3 was found across multiple bacterial clades (A, B, D, and E), suggesting that *Hamiltonella* genomic background may impose greater constraints on APSE host range than viral factors alone.

*Hamiltonella* clade A showed a strong positive association with APSE2 (74.2%, n = 132, Shannon H’ = 0.91), while clade B displayed the highest APSE diversity (H’ = 1.01, 5 types) with APSE8 being the most common associate (58.7%) (Figure 3B).

Toxin-APSE associations demonstrated even stronger constraints than *Hamiltonella*-APSE relationships, revealing characteristic patterns (Figure S2): APSE1 predominantly carried *stlx*, APSE2/8/9 variants typically harbored *cdtB*, and APSE3/11 were associated with *ydP*. Multinomial logistic regression predicting toxin from APSE variant showed excellent performance (pseudo R² = 0.63, accuracy = 89.8%, χ² = 755.9, df = 12), outperforming the clade- APSE model. All toxin types showed highly significant associations after FDR correction: *stlx* (χ² = 521.12, p < 0.001), *cdtB* (χ² = 421.24, p < 0.001), and *ydP* (χ² = 344.85, p < 0.001). The strongest predictive associations included APSE1 → *stlx* (OR = 617.90, 95% CI: 213.40- 1789.19) and APSE3 → *ydP* (OR = 76.92, 95% CI: 33.94-174.36), with near-perfect reciprocal specificity: APSE1 nearly exclusively encoded *stlx* (95.9%) while *stlx* was highly associated with APSE1 (89.9% of 158 isolates). Detailed analysis of *cdtB*-containing isolates revealed strong variant-specific associations: APSE2 exclusively carried *cdtB1* (41/41), APSE8 contained *cdtB2* (17/17), and APSE9 encoded *cdtB3* (3/3). In contrast, *ydP* showed the broadest APSE distribution (Shannon H’ = 1.24), indicating relaxed evolutionary constraints compared to other toxins.

The superior predictive performance of toxin-APSE associations (89.8% accuracy) versus clade- APSE associations (66.9% accuracy) suggests phage move horizontally among *Hamiltonella* more readily than toxins among APSE backbones, possibly due to functional compatibility requirements or co-evolutionary history.

Notably, despite high levels of diversity overall, just five *Hamiltonella*-APSE-toxin combinations (*A*-APSE2/*cdtB1*, *B*-APSE1/*stlx*, *B*-APSE8/*cdtB2*, *D*-APSE3/*ydP*, and *E*- APSE8/*cdtB2*) account for 95% of symbiont isolates recovered (Figure 4E, F, Figures S2 and S3). This concentration suggests that only a subset may be ecologically successful or evolutionarily stable across a given time and place.

**Figure 4:**
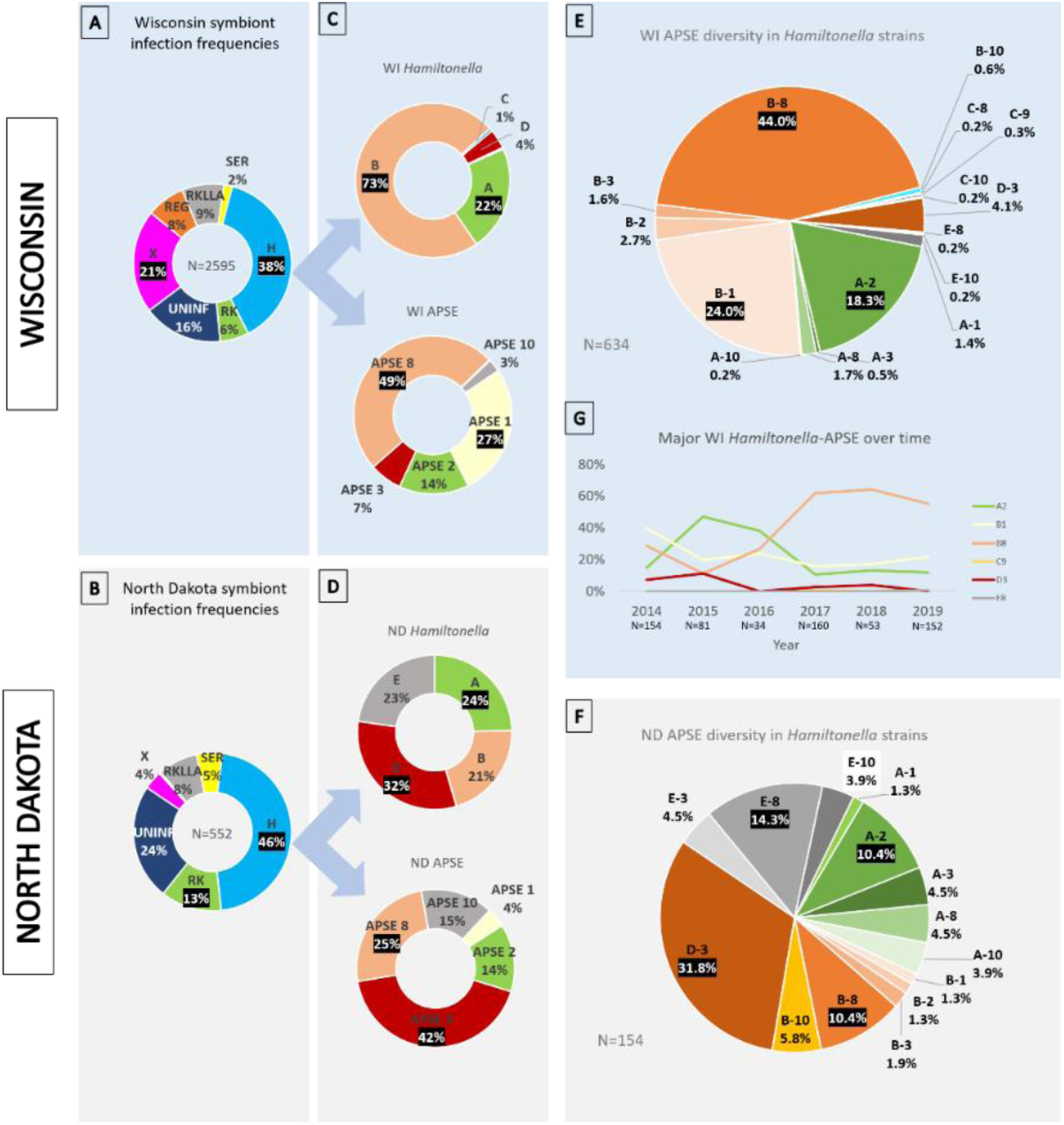
Temporal and Spatial Dynamics of *Hamiltonella* and Associated APSE Phage Variants in Pea Aphids. This figure compares prevalence and diversity across Wisconsin (WI) and North Dakota (ND). **Facultative Symbiont Infection Frequencies:** Relative frequencies of seven facultative symbionts and uninfected individual in pea aphid samples collected from **(A) Wisconsin** and **(B) North Dakota.** Distribution of bacterial clades and specific APSE variants within *Hamiltonella*-infected aphids from **(C) Wisconsin** and **(D) North Dakota**. Breakdown of *Hamiltonella* strains and associated APSE combinations across **(E) Wisconsin** and **(F) North Dakota**. Each slice represents a specific combination (e.g., B-8 is *Hamiltonella* clade B isolate with APSE8).**G. Temporal Dynamics in Wisconsin:** Changes in the relative frequencies of the major *Hamiltonella*-APSE combinations (A2, B1, B8, D3, E8) over the five-year sampling period (2014–2019) at the Wisconsin site. The number of samples (N) analyzed per year is indicated below the plot. *Hamiltonella* (H), *Rickettsia* (RK), *Fukatsuia* (X), *Rickettsiella* (RKLLA), *Regiella* (REG), *Serratia* (SER), No facultative symbionts (UNINF).

### Coinfections involving specific *Hamiltonella* strains

Fisher’s exact tests revealed highly significant, non-random associations between *Hamiltonella* clades and three common coinfecting symbionts, with distinct clade-specific preferences emerging for each symbiont type. For example, *Rickettsia* exhibited a strong preference for Clade A coinfections (Figure 3E, G: FDR-corrected χ² p < 0.001). Clade A showed 3.3-fold higher odds of *Rickettsia* coinfection compared to Clade B (OR = 3.30, 95% CI: 1.205-9.10, FDR-adjusted p = 0.012), while Clades D and E showed significantly reduced odds of association (OR = 0.22-0.23, both p < 0.001). *Rickettsiella* coinfections were more distributed across clades but still significantly non-random, with all clades showing reduced odds compared to Clade B (ORs: 0.13-0.31, all FDR-adjusted p < 0.01) (Figure 3I).

*Fukatsuia* was strongly associated with Clade B *Hamiltonella* (Figure 3F: FDR-corrected χ² p < 0.001), comprising 90% of all *Fukatsuia* associations, with Clade D representing the least preferred association (0.2%). Compared to Clade B, all other clades showed dramatically reduced odds of *Fukatsuia* coinfection: Clade A (OR = 0.101, 95% CI: 0.018-0.562, FDR-adjusted p = 0.004), Clade D (OR = 0.002, FDR-adjusted p < 0.00001), and Clade E (OR = 0.007, FDR-adjusted p < 0.00001).

This association between specific *Hamiltonella* strains and other symbiont species is strong enough that *Hamiltonella* strain diversity decreases overall in coinfections (Simpson’s index D = 0.71) versus single infections (D = 0.83) because most symbionts pair with B-clade strains.

We also found unexpected instances of two *Hamiltonella* isolates infecting the same individual aphid (A+B, N=64; B+E, N=10; A+E, N=5). To test whether observed *Hamiltonella* coinfections occurred at frequencies expected from random pairing based on clade abundance, we performed χ² goodness-of-fit tests with FDR multiple comparison corrections. The overall multinomial test revealed that the coinfection pattern deviated significantly from random expectation (χ² = 77.4, df = 5, p < 0.001). Individual comparisons showed that A+B coinfections occurred significantly more frequently than expected under random pairing (χ² = 43.1, FDR-corrected p < 0.001). While A+E and B+E coinfections occurred at frequencies slightly higher than expected (A+E: observed = 5, expected = 4.1; B+E: observed = 10, expected = 8.7), these deviations were modest and only marginally significant after FDR correction (p = 0.049 and p = 0.006, respectively). More strikingly, several expected combinations were completely absent despite substantial expected frequencies (A+D: expected = 6.5, observed = 0, p = 0.016 FDR-corrected; B+D: expected = 13.6, observed = 0, p < 0.001 FDR-corrected). Other observed combinations were only modestly enriched, marginally significant (A+E: χ² = 4.2, p = 0.048 FDR-corrected; B+E: χ² = 8.7, p = 0.006 FDR-corrected). These results suggest that the persistence of *Hamiltonella* strain coinfections may not be solely determined by clade abundance, but rather may reflect biological interactions.

We also examined correlations between APSE variants and the propensity of associated *Hamiltonella* strains to form coinfections with other symbiont species. Fisher’s exact tests revealed significant non-random associations between APSE variants and three of the four symbiont taxa examined: *Fukatsuia*, *Rickettsiella*, and *Rickettsia,* all of which showed highly significant associations (FDR-adjusted p < 0.0001) with specific APSE variants. In contrast, *Regiella* showed no significant association (FDR-adjusted p = 0.478). However, findings that specific APSEs strongly associate with particular *Hamiltonella* clades (Figure 3B) complicate interpretations of these comparisons. For example, it is unsurprising that *Rickettsia* is primarily associated with APSE2, given this symbiont’s significant association with A-clade *Hamiltonella* (Figure 3E) which, in turn, usually carries APSE2 (Figure 3B). Yet, since B-clade *Hamiltonella* are associated with diverse APSEs (Figure 3B), there is potential to ascertain the influence of APSEs on symbiont coinfections independent of *Hamiltonella* genotype. A chi-square test of independence confirmed a significant association between B-clade associated APSE variants and symbiont community composition (p < 0.0001). Fisher’s exact tests revealed highly significant non-random associations between specific phage variants in B-clade *Hamiltonella* and three commonly coinfecting symbionts: *Fukatsuia*, *Rickettsiella*, and *Rickettsia* (all FDR-adjusted p < 0.0001). Specifically, B1 and B8 were strongly associated with *Fukatsuia* (81.3% and 91.7%, respectively), B3 with *Rickettsiella* (66.7%), and B2 and B10 showed elevated associations with *Rickettsia* (37.5% and 42.9%, respectively).

### Symbiont and APSE Diversity in Wisconsin vs North Dakota Pea Aphid Populations

We conducted diversity analyses of symbiont communities and associated APSE phages from pea aphids collected on *Medicago* at Wisconsin and North Dakota sites (Figure 4). Analysis of 3,147 individual aphids revealed seven symbiont species at both sites, but with contrasting diversity patterns. Wisconsin exhibited modestly higher overall symbiont Shannon diversity (H’ = 1.639 vs 1.462; t = 4.749, p < 0.001), driven primarily by greater evenness (J’ = 0.842 vs 0.751). However, compositional differences were pronounced: *Hamiltonella* dominated both communities but was more prevalent in North Dakota (46.2% vs 38.5%), while *Fukatsuia* was substantially more abundant in Wisconsin (21.2% vs 3.8%). Aphids lacking facultative symbionts were more common in North Dakota (23.7% vs 16.3%).

Among 785 *Hamiltonella*-infected individuals, the pattern reversed dramatically (Figure 4C, D). North Dakota showed significantly higher *Hamiltonella* clade diversity (H’ = 1.373 vs 0.722; t = 18.29, p < 0.001) with near-perfect evenness (J’ = 0.990 vs 0.449). Isolate from the same five clades (A-E) were recovered, but their distributions differed strikingly between sites (χ² = 285.97, p < 0.001; Bray-Curtis dissimilarity = 0.75). North Dakota maintained balanced representation across clades D (31.8%), A (24.7%), E (22.7%), and B (20.8%), with clade C absent (but note the small sample size). Wisconsin was overwhelmingly dominated by clade B (73.2%), with all other clades comprising less than 25% of the community.

Analysis of 630 APSE-containing isolates revealed seven phage types in both locations, but North Dakota again showed higher diversity (Figure 4C, D: H’ = 1.464 vs 1.285; t = 2.70, p = 0.007) and evenness (J’ = 0.817 vs 0.660), with significant compositional differences (χ² = 105.50, p < 0.001; Bray-Curtis dissimilarity = 0.514). North Dakota displayed a more balanced APSE distribution, led by APSE1 (40.5%), APSE3 (25.9%), and APSE8 (17.1%), while Wisconsin was dominated by APSE8 (54.2%), with APSE1 (17.4%) and APSE2 (16.1%) as secondary types.

Seventeen distinct *Hamiltonella*-phage combinations were recovered (N = 789). North Dakota showed substantially higher combination diversity (Figure 4E, F) H’ = 2.236 vs 1.561; t = 7.71, p < 0.001) and evenness (J’ = 0.826 vs 0.563), with extreme compositional divergence (χ² = 367.15, p < 0.001; Bray-Curtis dissimilarity = 0.782). North Dakota maintained relatively balanced representation with D3 (31.4%), E8 (14.1%), and A2 (10.3%) as leading combinations. Wisconsin was overwhelmingly dominated by B8 (44.1%) and B1 (24.0%), which together comprised 68.1% of all combinations.

Chao1 richness estimation indicated very high sampling completeness (94.4%), with only one additional combination likely remaining undetected. The 17 observed combinations represent 48.6% of the theoretical maximum (35), suggesting that many possible Hamiltonella-APSE pairings either do not occur naturally or are absent from these populations.

In our analysis of APSE variant plus toxins (N = 626), we recovered 16 distinct combinations. Once again, North Dakota maintained higher combination diversity (Figure S3: H’ = 2.023 vs 1.554; t = 4.80, p < 0.001) and evenness (J’ = 0.814 vs 0.560), with significant compositional differences (χ² = 175.30, p < 0.001; Bray-Curtis dissimilarity = 0.728). North Dakota showed a balanced distribution led by *ydP*-APSE3 (28.7%), *cdtB*-APSE8 (19.8%), while Wisconsin was dominated by *cdtB*-APSE8 (45.9%) and *sltx*-APSE1 (26.1%), comprising 72% of all combinations.

### Temporal Variation

Analysis of annual variation at the better-sampled Wisconsin site (2014-2019) revealed substantial temporal dynamics in symbiont communities, with large fluctuations in facultative symbiont species and *Hamiltonella*-APSE combinations (Figure 4G. Four of seven symbiont categories showed significant temporal trends over the study period (Figure S4A). *Hamiltonella* significantly increased ( β = 0.085 ± 0.022 SE, χ² = 14.65, p < 0.001), with odds increasing by 8.9% per year (OR = 1.09). Conversely, three symbionts showed significant declining trends: *Regiella (*β = -0.220 ± 0.089 SE, χ² = 6.14, p = 0.013) with odds decreasing by 19.7% per year (OR = 0.80), *Rickettsiella* (β = -2.466 ± 0.294 SE, χ² > 50, p < 0.001) with odds decreasing by 91.5% per year (OR = 0.08), and *Serratia* (β = -0.452 ± 0.194 SE, χ² = 5.43, p = 0.020) with odds decreasing by 36.3% per year (OR = 0.64). *Fukatsuia* (χ² = 1.49, p = 0.223), *Rickettsia* (χ² = 1.49, p = 0.222), and uninfected aphids (χ² = 0.086, p = 0.769) showed no significant temporal trends.

Symbiont community diversity declined significantly over the study period. Shannon diversity, which emphasizes species richness, decreased (β = -0.063 ± 0.019 SE, F₁,₄ = 11.14, p = 0.028), as did Simpson diversity (β = -0.019 ± 0.005 SE, F₁,₄ = 14.07, p = 0.020), which emphasizes evenness. Correspondingly, community dominance increased (β = 0.017 ± 0.005 SE, F₁,₄ = 12.75, p = 0.021), indicating greater concentration of abundance among fewer symbiont types. Species richness (F₁,₄ = 0.58, p = 0.490) and evenness (F₁,₄ = 2.85, p = 0.133) showed no significant temporal trends.

Despite significant changes in individual symbionts and diversity metrics, PERMANOVA detected no significant overall compositional change (F₁,₄ = 1.45, R² = 0.27, p = 0.233), suggesting that the fundamental community structure remained relatively stable.

The Wisconsin symbiont community underwent significant simplification from 2014-2019, characterized by the virtual elimination of *Rickettsiella*, declines in *Regiella* and *Serratia*, and increasing dominance by *Hamiltonella* (from 35.1% to 46.2% of the community). This resulted in measurable decreases in Shannon and Simpson diversity indices, while overall community structure remained relatively stable as assessed by multivariate methods.

Three of the four common *Hamiltonella* + phage combinations showed significant temporal trends over the six-year study period (Figure 4G). B8 showed a significant increasing trend (β = 0.408 ± 0.143 SE, χ² = 8.13, p = 0.004), with odds increasing by 50% per year (OR = 1.50).

Conversely, B1 and D3 showed a significant decreasing trends (B1; β = -0.166 ± 0.084 SE, χ² = 3.93, p = 0.047: D3; β = -0.513 ± 0.176 SE, χ² = 8.44, p = 0.004), with odds declining by approximately 15% (OR = 0.85) and 40% (OR = 0.60) per year per year, respectively. Despite being numerically abundant, A2 showed no significant temporal trend (β = -0.236 ± 0.166 SE, χ² = 2.03, p = 0.154).

At the community level, richness showed no significant temporal trend (F₁,₄ = 0.016, p = 0.906). However, Shannon diversity showed a marginally significant declining trend over time (β = - 0.071 ± 0.030 SE, F₁,₄ = 5.66, p = 0.076), suggesting potential simplification of the *Hamiltonella* + phage community structure. These results demonstrate significant temporal dynamics in *Hamiltonella* + phage associations within Wisconsin populations, with B8 increasing in prevalence while B1 and D3 declined over the study period. The marginal decline in Shannon diversity suggests overall community simplification, potentially driven by the expansion of B8 and contraction of other combinations.

## Discussion

Our survey of over 3,100 North American pea aphids reveals that *Hamiltonella defensa*– together with its APSE phages–is a highly dynamic and structurally central component of the heritable microbiome of pea aphids. By integrating strain-level typing with toxin and phage diversity analyses, we conservatively uncovered 38 distinct *Hamiltonella*–APSE–toxin combinations, substantially exceeding the diversity reported to date. This cryptic diversity highlights the modularity of the system, in which bacteriophage backbones and toxin cassettes are shuffled across bacterial lineages, generating novel phenotypes with the potential to alter parasitoid resistance and other phenotypes. The strain-level diversity observed for *Hamiltonella* does not appear to be the norm for other aphid facultative symbionts (Henry et al. 2013, Badji et al. 2021, Patel et al. 2023, Peng et al. 2023), making this symbiont particularly notable for understanding how modular evolution can generate functional variation.

Our findings demonstrate that strain-level variation likely drives both protective phenotypes and symbiont community assembly, with *Hamiltonella* acting as a central architectural element. Below, we explore the mechanisms generating this diversity, the ecological consequences of strain-specific interactions, and the implications for understanding defensive symbiosis and biological control.

### Symbiont Community Structure

Our analyses revealed a complex but highly structured symbiont community consistent with previous surveys (e.g., (Russell et al. 2013, Smith et al. 2015, Rock et al. 2018)), but extending these with comprehensive strain-level variation for the focal symbiont *Hamiltonella*. A large majority (75%) of *Medicago*-associated aphids harbored one or more facultative symbionts, with an average of 1.6 symbionts per individual for those carrying facultative symbionts.

*Hamiltonella* was the most prevalent facultative symbiont, infecting 56% of surveyed aphids, and exhibited a significant tendency toward coinfection, participating in 92% of the 35 observed species-level combinations (Table 1).

*Hamiltonella* + *Fukatsuia* pairing occurred 1.63 times more frequently than expected by chance, while other combinations--notably *Hamiltonella* + *Serratia*--were completely absent despite both symbionts being present in populations. While aphids with single symbionts receive specific benefits (Oliver et al. 2014), coinfections can confer multiple protective phenotypes, though outcomes vary with genotypic interactions (Oliver et al. 2006, Doremus and Oliver 2017, McLean et al. 2020, Weldon et al. 2020). Higher vertical transmission rates for enriched combinations and lower for depleted ones may create or reinforce these patterns (Rock et al. 2018). *Serratia* + *Rickettsiella* also showed significant enrichment in our study (Table 1), and metabolic complementarity, particularly in biotin synthesis, may facilitate their cooperation (Peng et al. 2023). Conversely, the *Hamiltonella* + *Serratia* combination, which can impose severe fitness costs (Oliver et al. 2006), was never observed, suggesting active selection against incompatible pairings. These patterns demonstrate that symbiont community assembly is not purely a stochastic process, but is shaped by host-level natural selection and a complex array of positive and negative interactions among community members.

### Modular Evolution and Horizontal Transfer

The recurrent horizontal transfer of APSE phages among *Hamiltonella* lineages, and toxin cassettes among phage backbones, creates a combinatorial defense system. Chao1 estimates suggest 94% sampling completeness, with the 17 observed types representing only about 50% of the theoretical possibilities. Whether absent combinations are biologically unstable or simply rare/transient remains unknown. This modular architecture enables evolutionary innovation while maintaining functional stability, potentially allowing aphids to rapidly track parasitoid adaptation (Degnan and Moran 2008, Rouïl et al. 2020, Boyd et al. 2021, Patel et al. 2023).

However, our predictive modeling reveals asymmetric constraints on this modularity: APSE backbones and toxin identity show strong correspondence (89.8% predictive accuracy), while *Hamiltonella* clades and APSE variants exhibit looser associations (66.9% accuracy). This pattern indicates that APSE phages move more readily among bacterial lineages (e.g., APSE3 occurring in four *Hamiltonella* clades and five APSE variants within B-clade alone) than toxin modules among phage backbones. Such asymmetry likely reflects different evolutionary constraints: While modular recombination is an important source of evolutionary novelty, most of these events are expected to create dysfunctional phages, which are rapidly removed from the population (Hendrix et al. 1999, Hatfull 2008). For example, modular recombination may produce regulatory circuits that do not interact properly, leading to poor assembly, failed replication, or other outcomes. The near-perfect associations observed between specific APSE variants and specific toxins – i.e. APSE1 carrying *sltx* (96%), APSE2 exclusively encoding *cdtB1*, and APSE3 carrying *ydP* – further support strong functional constraints on toxin-phage compatibility.

In contrast, intact phage transfer between *Hamiltonella* isolates is likely constrained primarily by ecological opportunities. These include opportunities for phage particles to move laterally through plant phloem or wasp ovipositors to establish in new bacterial genotypes.

Horizontal transfer of their host *Hamiltonella* cells could occur by similar mechanisms (Caspi-Fluger et al. 2012, Gehrer and Vorburger 2012, Pons et al. 2019), facilitating multiple *Hamiltonella* strain infection and an additional means for phage transfer between these symbiont lineages. In the present work, we found several coinfections involving two *Hamiltonella* strains, which provide opportunities for the swapping of phage virulence modules or toxins within modules.

Some *Hamiltonella* strains lack APSEs (e.g., (Degnan and Moran 2008) and others are associated with the repeated spontaneous loss of APSE (Oliver and Moran 2009, Weldon et al. 2013). Some *Hamiltonella* cells within APSE-harboring aphids potentially lack integrated prophage, created a reservoir of cells where APSE infection can establish. Each of these may create conditions that promote the establishment of APSE. Conversely, some APSE variants may not produce infective particles, which may limit transfer potential.

Regardless of whether phage acquisition stems from swapping or infection of uninfected cells, our findings here suggest *Hamiltonella* vary in their abilities to accept novel phage lineages. Most hospitable is the B clade with five APSE variants. Least hospitable was the D clade, where 100% of isolates carried APSE3 (N = 89). Ironically, despite this apparent specificity, most phage loss events in the lab involved D3 *Hamiltonella* losing APSE3 (Oliver and Moran 2009, Weldon et al. 2013), suggesting that even seemingly stable associations can be fragile.

### Community Architecture: *Hamiltonella* as a Central Hub

We found that *Hamiltonella* strains vary not only in their defensive repertoire but also in their propensity to form coinfections with other symbionts. B-clade *Hamiltonella* disproportionately co-occurs with *Fukatsuia* (90% of *Fukatsuia* coinfections) and *Rickettsiella*, while A-clade strains associate primarily with *Rickettsia* (Figure 3). These non-random associations suggest that *Hamiltonella* acts as a central partner structuring the broader symbiont community, with *Hamiltonella* strain identity influencing which other symbionts successfully colonize or persist.

Two non-exclusive mechanisms may underlie these patterns. First, metabolic complementarity or co-dependency: the near-obligate nature of *Fukatsuia* coinfection (95% coinfected, 97% with *Hamiltonella*) suggests possible dependencies, perhaps involving biosynthetic pathways where genome-degraded symbionts rely on partners to compensate for lost functions—a pattern documented for *Serratia*-*Rickettsiella* interactions in biotin synthesis (Peng et al. 2023). Second, mobile element-mediated effects: since *Hamiltonella* clades differ primarily in their complement of insertion sequences and prophages (Degnan et al. 2009, Chevignon et al. 2018, Patel et al. 2023), these elements may modulate host immunity or bacteriocyte physiology in ways that differentially affect other symbionts. The observation that specific APSE variants within B-clade show distinct coinfection preferences (B1/B8 with *Fukatsuia*, B3 with *Rickettsiella*) further implies that phages themselves may shape community assembly, either by altering host physiology or by mediating microbe-microbe interaction; a possibility requiring further investigation.

Together, these results suggest either within-aphid ecology, with cooperation or competition among coinfections, or host-level selection, where successful phenotypes may emerge not from single infections but from multi-symbiont partnerships (Rock et al. 2018, Weldon et al. 2020, Carpenter et al. 2021, Peng et al. 2023). The finding that certain *Hamiltonella* coinfections (A+B) occur significantly more frequently than expected by chance, while others (A+D, B+D) are completely absent despite substantial expected frequencies, further supports the hypothesis that biological interactions drive observed patterns of symbiont distributions.

### Temporal and Geographic Dynamics

Our results align with prior findings of geographic differences in pea aphid symbiont communities and the rapid turnover of facultative symbionts over time (Russell et al. 2013, Smith et al. 2015, Ives et al. 2020). Further, our strain-level resolution reveals patterns that challenge simple predictions based on laboratory-measured parasitoid resistance. In our study, we found North Dakota pea aphid populations exhibited significantly higher *Hamiltonella* and APSE diversity than Wisconsin, with balanced representation across lineages. In contrast, Wisconsin populations were dominated by B-clade, which rose from 32% to 80% of *Hamiltonella* isolates sampled over five years.

Our strain-level resolution reveals that this cryptic diversity is subject to strong and fluctuating selection. The different strain compositions between these populations suggest local adaptation to regional parasitoid communities or environmental conditions (Higashi et al. 2020), consistent with aphid studies showing that *Hamiltonella* genotypes correlate with local parasitoid pressures (Wu et al. 2022, Wu et al. 2025). However, the temporal dynamics in Wisconsin reveal a striking paradox: the most common *Hamiltonella*/APSE over time in Wisconsin are those associated with conferring intermediate levels of resistance to *A. ervi* parasitoids in laboratory assays (Oliver and Higashi 2019): A2 early (Figure 4, 2015-16), later replaced by B8 (Figure 4, 2017-2019).

Meanwhile, D3, the most protective combination in laboratory bioassays (Moran et al. 2005, Martinez et al. 2014, Martinez et al. 2018a), declined to near-absence despite field evidence of effectiveness (Ives et al. 2020). This suggests that ecological success is not dictated by parasitoid resistance alone, but by trade-offs involving transmission efficiency (Weldon et al. 2013), metabolic costs, or compatibility with beneficial coinfections. Indeed, D-clade *Hamiltonella* rarely participated in coinfections (<5%), while B-clade was involved in 92%. The dominance of B1 and B8, which confer relatively weak protection in bioassays, suggests that multi-symbiont strategies or alternative fitness components may outweigh maximal parasitoid resistance under field conditions (Smith et al. 2021).

### Implications and Future Directions

These findings complicate biological control strategies targeting aphid pests. Parasitoid introductions may initially succeed against prevalent symbiont strains, only to face rapid replacement by pre-existing, resistant variants, a process that occurs within or between seasons rather than over decades. Our documentation of extensive standing variation means that aphid populations possess substantial capacity for evolutionary rescue, potentially undermining control efforts. Our findings further indicate that introduced pest species may carry or create ample variation in defensive symbionts to thwart biological control effects (Desneux et al. 2018).

Predicting the stability of biological control, therefore, requires strain-level monitoring rather than species-level symbiont surveys. For the pea aphid system, genotyping of anti-parasitoid APSE will be a key part of such assessment.

More broadly, the bacterial-phage-toxin combination creates a modular system resembling eukaryotic immune receptors where diversity arises through recombination (Hafer-Hahmann and Vorburger 2021). The structured nature of coinfections, where certain combinations preferentially co-occur while others are mutually exclusive, further implies that heritable microbiomes function as integrated consortia rather than collections of independent agents.

Beyond parasitoid defense, this standing variation in symbiont genotypes may also provide aphids with adaptive capacity to respond to climate change and other anthropogenic stressors, extending the implications of defensive symbiont diversity to broader ecological challenges (Russell and Moran 2006, Doremus et al. 2018, Higashi et al. 2020, Oliver and Higashi 2021, Tougeron and Iltis 2022).

In sum, our findings demonstrate that *Hamiltonella* and its phages function as a modular and rapidly evolving defensive system embedded within a wider symbiont community. Strain-level variation drives both protective phenotypes and community assembly, producing complex, shifting associations that enable aphid populations to contend with parasitoid attack and other ecological challenges. Recognizing this hidden diversity emphasizes the need for strain- and phage-level resolution in symbiosis research, with implications for understanding host-parasite coevolution, predicting ecosystem responses to biological control, and appreciating how microbial consortia—rather than individual symbionts—shape host adaptation.

## Acknowledgements

We thank UGA undergraduate researchers Alexandria Maddox, Catrina Chamberlain, Collin Pannell, Dorbin Abendano, and Nhu-Y Phan for technical assistance. This was supported by NSF Awards 1240892 and 2240392 to KMO and 1050098 and 2240393 to JAR.

