## Supplementary_materials for "Modular evolution and strain-specific partnerships: how *Hamiltonella defensa* shapes defense and symbiont communities in aphids"

### Table S1: Primers, reaction conditions, and references used.

| Primer names | Target | Product | Sequence 5' to 3' Forward primer | Reverse primer | Length (bp) | Credit |
|---|---|---|---|---|---|---|
| ***H. defensa* PCR diagnostics** | | | | | | |
| NY26_F1<br>NY26_R1 | *H. defensa* strain A | RTX toxin bound to IS630 | TTTTTGGGGCGCAGACATTG | GTCACTCAGCTGCTCCTACC | 914 | Germain Chevignon (ref?) |
| PF1<br>PR1 | *H. defensa* strain B | Beta-lactamase class C bound to IS3 ssgr IS3 | CTCACACCCATGGCTTCACT | AGCTTTCAGCCATCTGCCAT | 780 | This paper |
| CCF1<br>CCR1 | *H. defensa* strains C | T1SS secreted agglutinin RTX CDS | GGACTCTAAAGATCAAGAAGGCA | ATTTAGTGATGTTGGGCGGGA | 500, 250 | This paper |
| ?? | *H. defensa* strains C, B | T1SS secreted agglutinin RTX CDS | GGACTCTAAAGATCAAGAAGGCA | ATTTAGTGATGTTGGGCGGGA | 500, 250 | Jacob/Melissa? |
| NY26_F1<br>NY26_R1 | *H. defensa* strain D | RTX toxin bound to IS630 | TTTTTGGGGCGCAGACATTG | GTCACTCAGCTGCTCCTACC | 500 | Germain Chevignon |
| YF6<br>YR6 | *H. defensa* strain D | Hypothetical protein bound to IS630 | CCTTGGGATGACGCAGAGTT | GTCTTTTGCGGGTTGTCCAC | 503 | This paper |
| ZA17_qPCR_F1<br>ZA17_R1 | *H. defensa* strain E | IS6 family transposase bound to transcriptional regulator | GCCCTGTTAAATGGCGTGTA | AATTGAGCAAGTGAAGCGTA | 573 | Germain Chevignon |
| ZA17_F1<br>ZA17_qPCR_R1 | *H. defensa* strain E | IS6 family transposase bound to transcriptional regulator | TACGTACTGTGATGCGCTGG | TGATTCTATTTAATTACCAGAGCGA | 500 | Germain Chevignon |
| ***H. defensa* sequencing diagnostics** | | | | | | |
| recJ44F<br>recJ1012R | *H. defensa recJ* gene | 5' –> 3' exonuclease | ATCCGCTCTCAGAAACATACC | GATGACATAAATCCAATGCCTC | 930 | Degnan and Moran 2008 |
| accD291F<br>accD832R | *H. defensa* accD gene | Carboxyl transferase, subunit β | TTCTGGAGCACAAAAAGACAC | AAGGTTCAGGTTGATGAGTCAG | 500 | Degnan and Moran 2008 |
| murE16F<br>murE936R | *H. defensa murE* gene | UDP-N-acetylmuramoylalanyl-Dglutamate 2,6-diaminopimelate | ACTAACGGGAAAACCACTAATAC | TTGAGAATGTCAGCGGTAATC | 1040 | Degnan and Moran 2008 |
| ptsI181F<br>ptsI709R | *H. defensa ptsI* gene | Phosphotransferase enzyme I | ATTTTACGGGCTTCTGCTTTTG | CTTCGGTGGTTGATTGACTCAG | 500 | Degnan and Moran 2008 |
| hrpA106F<br>hrpA984R | *H. defensa hrpA* gene | Predicted ATP-dependent helicase, involved in mRNA processing | AAACCCAATCTGACAAAAATAGG | TAACTCTTCGGCTTCTGACAAC | 860 | Degnan and Moran 2008 |
| **APSE sequencing diagnostics** | | | | | | |
| APSE1.1F<br>APSE2.4R | APSE P3 gene | Predicted virulence-associated protein | TCGGGCGTAGTGTTAATGAC | TTCCATAGCGGAATCAAAGG | | Degnan and Moran 2008 |
| newP3 SET 1 FWD<br>newP3 SET 3 REV | APSE P3 gene | Predicted virulence-associated protein | TTATTCCGTCGGGCGTAGTG | CGATTGCCCATATCGACCCA | 710 | This paper |
| NewP35 1F<br>NewP35 2R | APSE P35 gene | Phage DNA transfer protein | CTAGCTGGGGAAACAGTGGG | GTTGCGAGCGATGCGTTTAT | 810 | This paper |
| P38FWD1<br>P38REV1 | APSE P38 gene | Phage integrase | CGAAGCCTGCTTGAGATGGA | CGCGTCAGAATACAGCAAGC | 550 | This paper |
| APSE30.6F<br>APSE31.9R | APSE P45 gene | DNA polymerase | ACGGCACTTAAACGCTATCC | TGGGATGTGTATGGACGTTG | 600 | Degnan and Moran 2008 |
| **Secondary sequencing diagnostics** | | | | | | |
| RECJ FWD 1<br>RECJ REV 1 | *H. defensa recJ* gene | 5' –> 3' exonuclease | GCGCGAGGGACAAAAAGTTT | AGGGTAGGCTCGGTCAATCT | 1100 | This paper |
| ACCDF1<br>ACCDR1 | *H. defensa* accD gene | Carboxyl transferase, subunit β | CGCGGTTACATGCCTTCTTG | ACTGGCTAACTTGTGTCGCA | 650 | This paper |
| MUREF1<br>MURER1 | *H. defensa murE* gene | UDP-N-acetylmuramoylalanyl-Dglutamate 2,6-diaminopimelate | CCATCCTGGCTGAAGCAGAA | TGTTTCAGCGCATCAGGAGT | 900 | This paper |
| PTSIF1<br>PTSIR1 | *H. defensa ptsI* gene | Phosphotransferase enzyme I | GGCTTGCCTGTCACTGATCT | GCAAACAGGCCGCACTTTTA | 1100 | This paper |
| HRPA FWD 1<br>HRPA REV 1 | *H. defensa hrpA* gene | Predicted ATP-dependent helicase, involved in mRNA processing | CGACGGCATTTTACTGGCTG | TGGGATATCGGCTCCAAGGA | 650 | This paper |
| **Toxin Sequence primers** | | | | | | |
| cdtBtox1F<br>cdtBtox1R | *cdtB* 1 & 2 | Cytolethal distending toxin subunit B | TGGCAATTCAGGAAGCTGGA | GCTATTCAGCTCGCTGGGAA | 450 | This paper |
| cdtBtox1BF<br>cdtBtox1BR | *cdtB* 1 & 2 | Cytolethal distending toxin subunit B | CACTCCGCCAATGCAAACAT | TTATTGCGCTGACATGGGGA | 100 | This paper |
| cdtBtox3F<br>cdtBtox2R | *cdtB 3* | Cytolethal distending toxin subunit B | CGGGATAGTCCTTGCGTGTT | CCTCGTTGCGGGTATCAACT | 680 | This paper |
| stx1F<br>stx2R | *stx* | shiga toxin alpha subunit | TGCTTGTTTTAGGCGGAGGT | GATCGGCGATGGTGGGTAAT | 940 | This paper |
| YD1F | YD | YD-repeat toxin | TGGGACTCGCAGTTTTCTCC | GGTTCGGACCACGTCTCTTT | 740 | This paper |

Figure S1A: Maximum likelihood tree (left) of five concatenated housekeeping loci and partitioned models. Neighbor-Net algorithm network analysis (right).

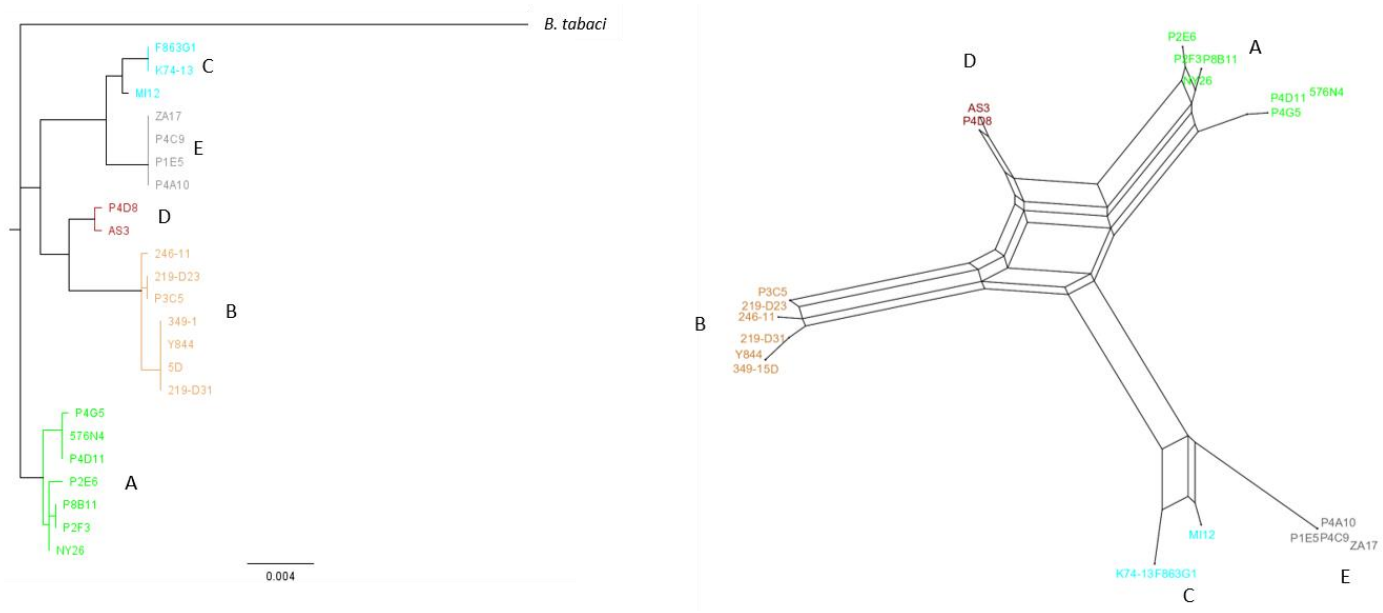

Figure S1B: Neighbor-Net algorithm network analyses of individual *Hamiltonella* housekeeping genes

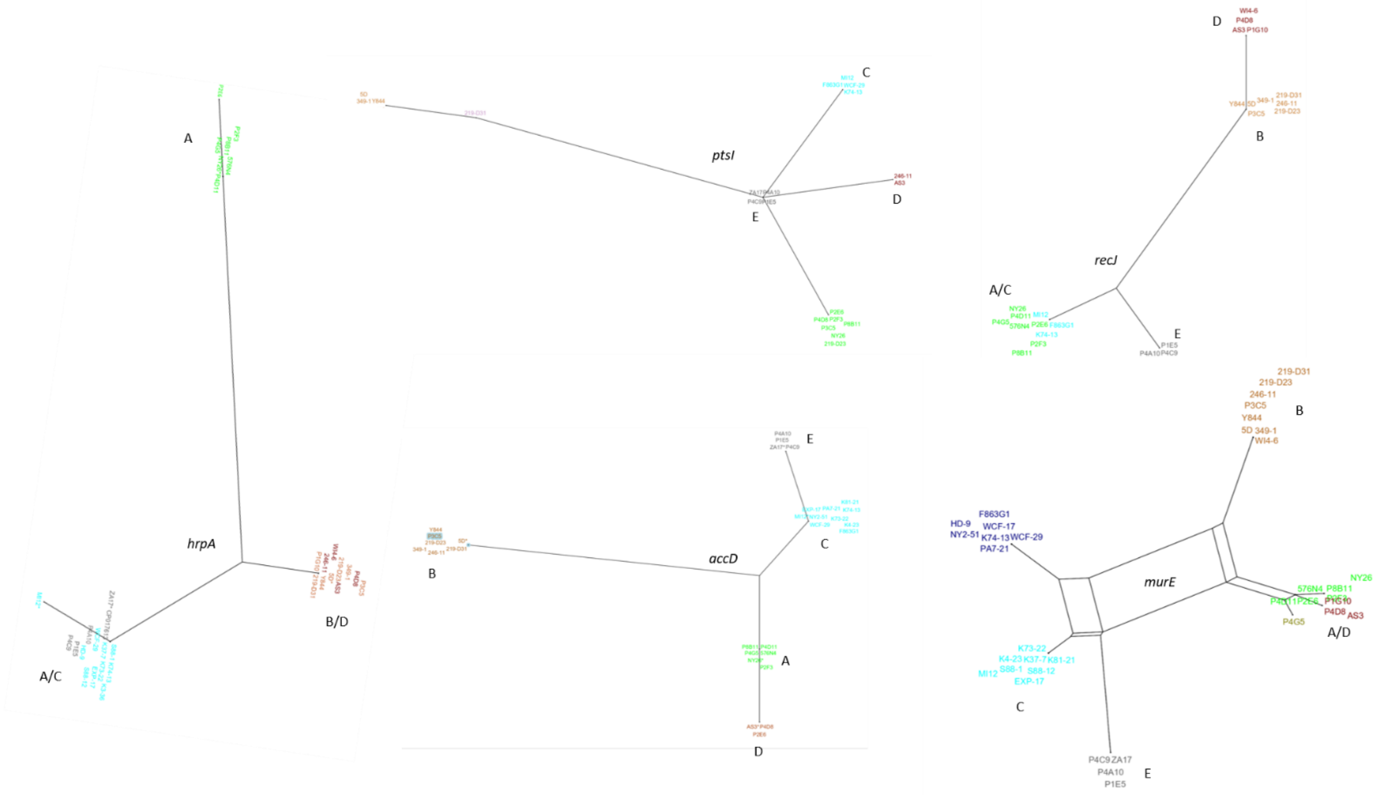


.

Figure S1C: Neighbor-Net algorithm network analyses of individual APE genes representing phage backbone modules

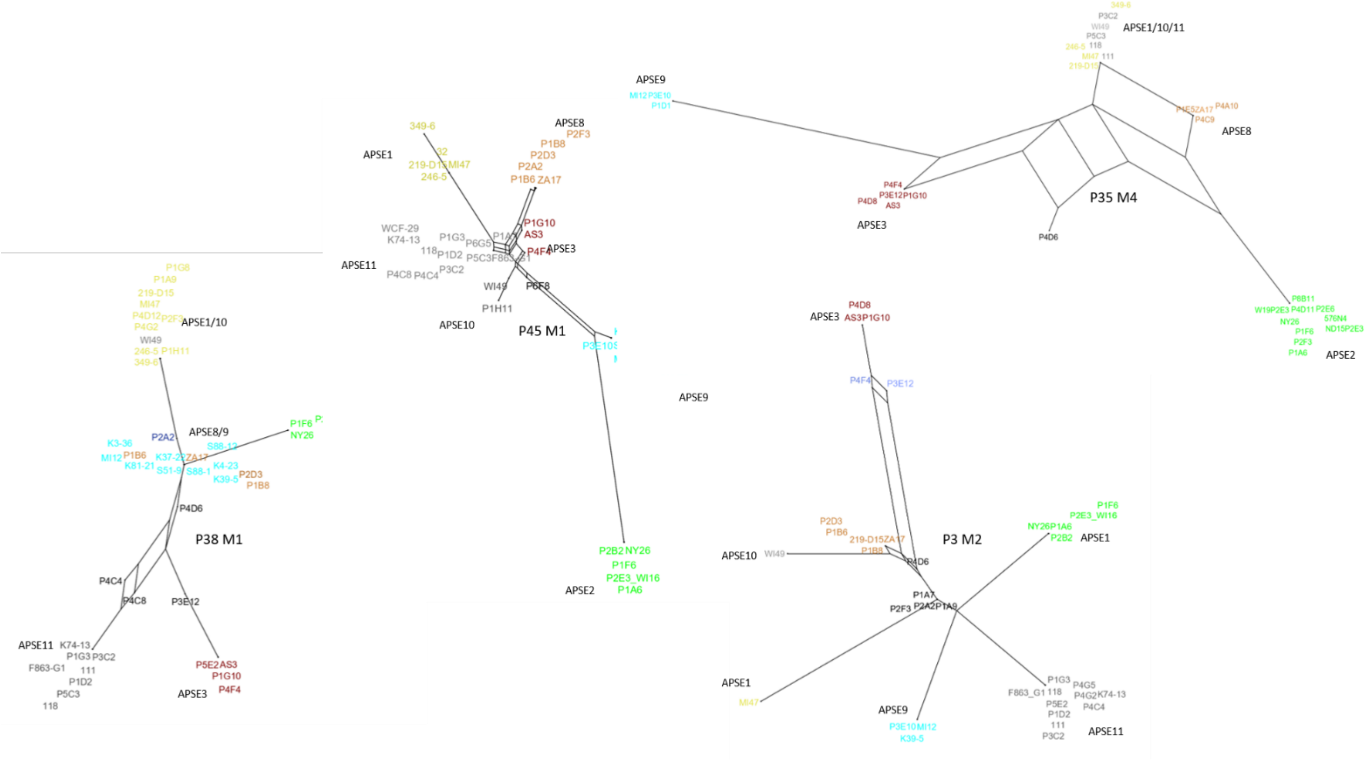

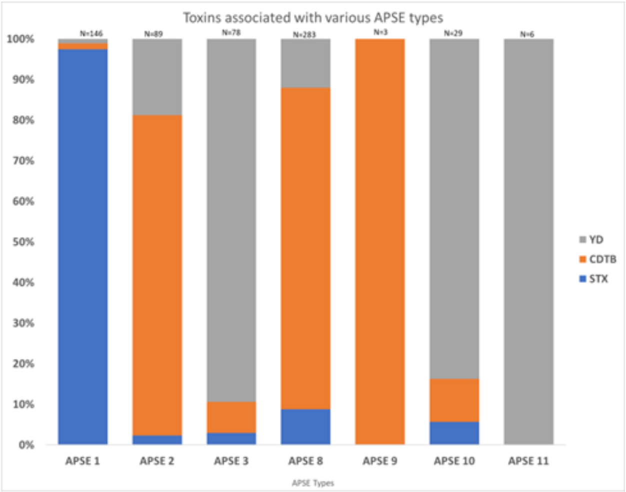


Figure S2: Proportion of APSE variants carrying either YDp, CDTB or SLTX toxins

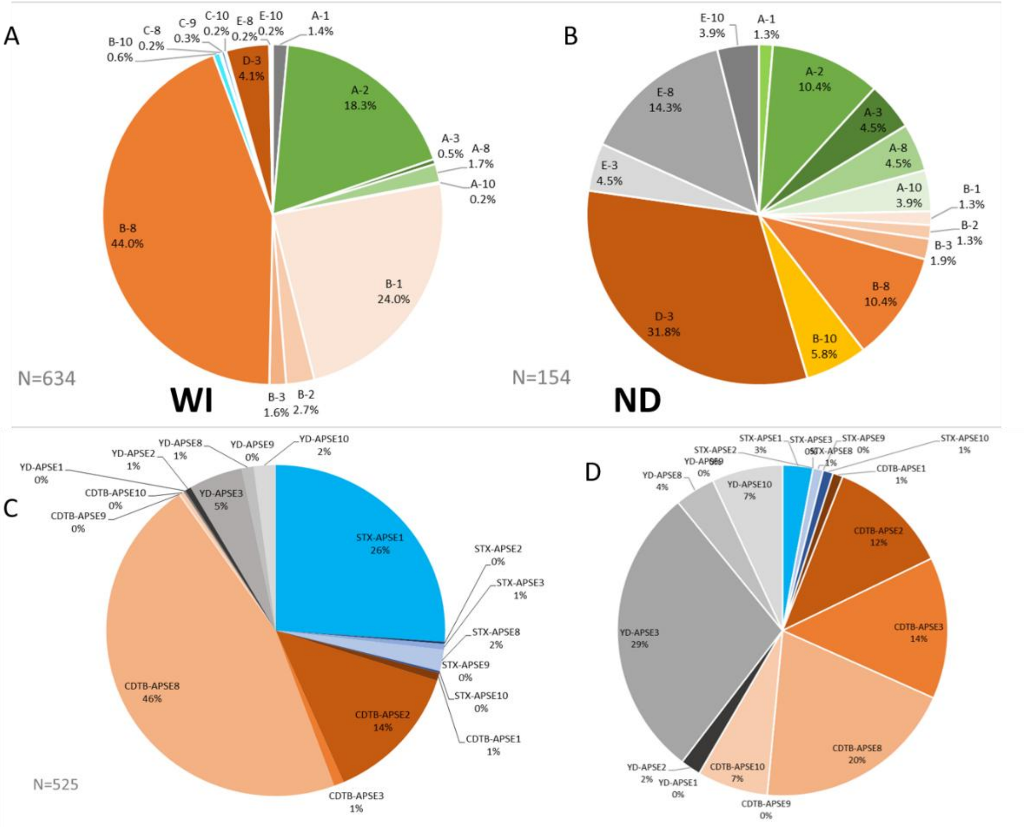


Figures S3. *Hamiltonella*-APSE (A, B) and APSE-toxin (C, D) combinations in WI (Wisconsin) and ND (North Dakota) pea aphid populations.

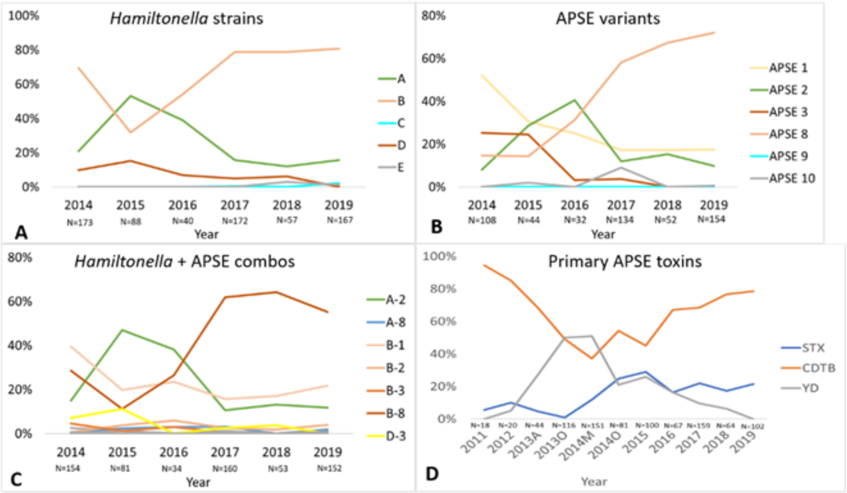


Figure S4. Temporal trends in symbiont species (A), APSE variants (B), *Hamiltonella*-APSE combinations (C), and phage-encoded toxins (D).
